# Neural correlates of future volitional action in *Drosophila*

**DOI:** 10.1101/2023.09.08.556917

**Authors:** Luke E. Brezovec, Andrew B. Berger, Shaul Druckmann, Thomas R. Clandinin

**Affiliations:** Fairchild D200, 299 W. Campus Drive, Department of Neurobiology, Stanford University, Stanford, CA, 94305

## Abstract

The ability to act voluntarily is fundamental to animal behavior^1,2,3,4,5^. For example, self-directed movements are critical to exploration, particularly in the absence of external sensory signals that could shape a trajectory. However, how neural networks might plan future changes in direction in the absence of salient sensory cues is unknown. Here we use volumetric two-photon imaging to map neural activity associated with walking across the entire brain of the fruit fly *Drosophila*, register these signals across animals with micron precision, and generate a dataset of ∼20 billion neural measurements across thousands of bouts of voluntary movements. We define spatially clustered neural signals selectively associated with changes in forward and angular velocity, and reveal that turning is associated with widespread asymmetric activity between brain hemispheres. Strikingly, this asymmetry in interhemispheric dynamics emerges more than 10 seconds before a turn within a specific brain region associated with motor control, the Inferior Posterior Slope (IPS). This early, local difference in neural activity predicts the direction of future turns on a trial-by-trial basis, revealing long-term motor planning. As the direction of each turn is neither trained, nor guided by external sensory cues, it must be internally determined. We therefore propose that this pre-motor center contains a neural substrate of volitional action.

## Main

Previous studies have identified neural signatures of internally generated actions in many species. In humans^6,7,8,9,10,11^, non-human primates^12,13,14,15^, cats^16^, rodents^17,18,19^, and zebrafish^20^ particular brain regions display an increase in activity, often referred to as the readiness potential, several seconds before initiating a future voluntary action. This phenomenon appears to emerge from spontaneous fluctuations in neural activity that can reflect planning and preparation of the movement^21^. However, these studies have relied on explicit tasks that constrain the timing or structure of behavioral actions, and it is unknown whether cognitive signals akin to the readiness potential exist in invertebrates.

Here, we take advantage of the fact that fruit flies spontaneously explore their environment, creating an opportunity to examine how specific actions are selected under task-free behavioral choices. Fly walking trajectories are highly structured, with saccadic turns interspersed with straight or curved runs^22,23,24,25,26,27,28^.

Specific neurons involved in turns versus forward movements have been identified, and many of the sensory circuits that guide movement have been described^29,30,31,32,33,34,35,36,37,38,39,40,41^. However, absent specific sensory stimuli that might bias the animal toward a particular trajectory, the neural mechanisms that underpin how the fly elects to turn in one direction or the other are unknown.

We sought to map neural activity associated with walking behavior across the fly brain and to quantitatively compare these signals across individuals. To do this we developed a pipeline for measuring neural activity across the whole volume of the brain while recording the animal’s locomotion in the dark (Fig. 1a)^42^. We expressed two fluorescent indicators in all neurons: GCaMP6f to monitor neural activity, and myristylated-tdTomato to capture brain structure^43,44^. Flies were head-fixed and the posterior head cuticle was removed to expose the brain (Extended Data Fig. 1). We then employed two-photon imaging with resonant scanning to achieve a volume imaging rate of 1.8 Hz, collecting 256 × 128 x 49 (x,y,z) voxels per volume, each occupying 2.6 × 2.6 × 5 μm. During each 30 minute imaging session the animal’s walking trajectory was recorded at 50Hz (Methods). At the end of each recording, we collected an anatomical scan, and registered every voxel of neural activity across individuals with high spatial accuracy (Methods; Extended Data Fig. 1)^45^. By registering all data into a common space and concatenating the neural and behavioral signals across flies, we created a unified dataset where 1.6M voxels have each been sampled 30,456 times over the course of 4.5 hr and 9 individuals (Fig. 1a). Finally, we reduced the number of features in the neural activity dataset by agglomerative clustering of neighboring voxels with similar responses to create superclusters (Fig. 1a; Methods). Such superclusters formed the basis for all subsequent analyses.

**Fig. 1:**
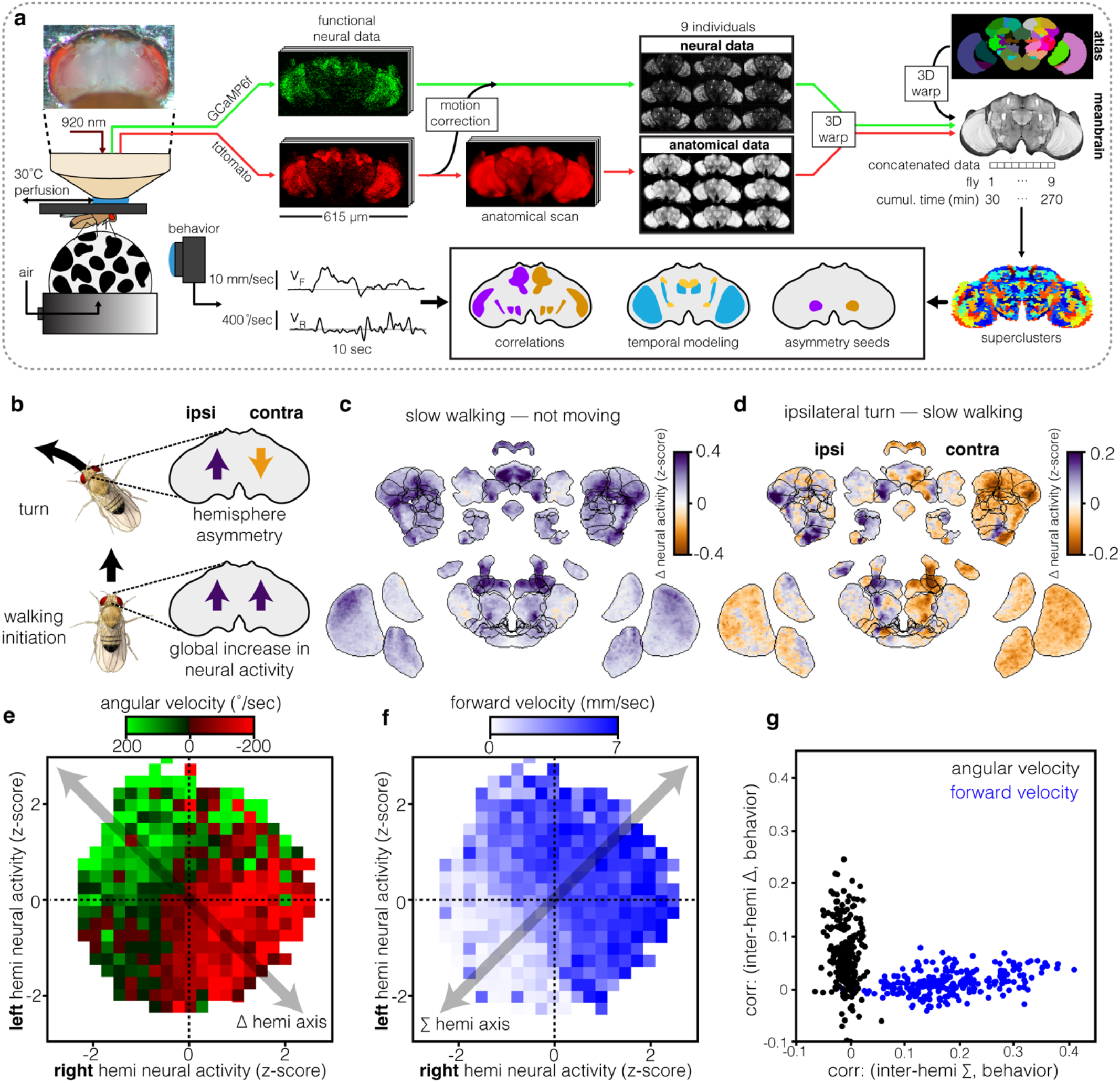
Whole-brain imaging in walking *Drosophila* reveals that neural representations of angular and translational velocity are widespread across the brain and orthogonal. **a**, Overview of the imaging pipeline. After dissection of the posterior head cuticle, flies were mounted under a two-photon microscope and walked on an air-suspended ball in complete darkness. GCaMP6f is expressed pan-neuronally, as is a structural marker, tdTomato. Brain volumes were acquired at 1.8 Hz to capture neural activity, while a subsequent anatomical scan was taken at higher spatial resolution. Nine individuals passed all quality control metrics (see Extended Data Fig. 1) and were used for subsequent analyses. All nine datasets were registered into a single mean brain using the structural marker; these warp parameters were then applied to the functional neural data to bring all flies into the same space. A standard anatomical atlas was also registered into this common space. Concatenating all data resulted in a “Superfly” with 270 cumulative minutes of neural and behavioral recording. Voxels were clustered into superclusters (Methods). Walking trajectories were decomposed into forward and rotational velocities. Data was analyzed using correlations, temporal modeling, and early hemisphere asymmetry analysis. **b**, Cartoon displaying neural activity changes associated with walking initiation and turning. The transition from not moving to moving is associated with a global increase in neural activity, while a turn is associated with an increase in ipsilateral activity and decrease in contralateral activity. **c**, “Exploded” brain with anatomically defined neuropiles separated to allow visualization of entire volume. The brain is colored by the difference in neural activity between two categorical behavioral conditions: not moving (<1mm/sec forward and <20deg/sec rotation) and slow straight walking (1< and >4 mm/sec forward and <20 deg/sec rotation). A maximum intensity projection through each neuropile is shown. **d**, Exploded fly brain colored by difference in neural activity between two categorical behavioral conditions: slow straight walking and ipsilateral turns. **e-g**, Walking behavior conditioned on neural activity reveals that angular velocity is encoded by the difference in neural activity between matched pairs of superclusters in each hemisphere, while translational velocity is represented by the summed signals across these pairs. **e**, Example supercluster within the Inferior Posterior Slope (IPS) that responds to both angular and translational velocity. Angular velocity conditioned on joint neural activity across the two hemispheres is shown. 90^th^ percentile of behavior in each bin is shown. **f**, As in **e**, but displaying forward velocity. **g**, Correlation between the interhemispheric difference in neural activity and angular velocity for each matched pair of superclusters (black dots), as well as the interhemispheric sum and translational velocity for each matched pair of superclusters (blue dots).

### Turns involve a wide-spread symmetry break across brain hemispheres

In the absence of any spatially patterned sensory cues, tethered flies spontaneously initiated bouts of walking activity composed of sequences of turns and straight runs (spanning seconds), separated by periods of quiescence and grooming (lasting tens of seconds). We reasoned that, under these conditions, neural activity might reveal signals that initiate movement and execute specific turning maneuvers. Consistent with this, we observed significant correlations between neural activity signals and behavior that were absent from the anatomical signals (Extended Data Fig. 2). Moreover, we could predict translational and rotational velocities across all flies using the principal components of neural activity (Extended Data Fig. 3).

To obtain an initial description of the neural correlates of behavior, we compared time-averaged neural signals when the fly was stationary, walking forward in a straight line, and turning (Fig. 1b-d). Consistent with previous work, we observed a global increase in neural activity across the brain upon the initiation of movement (Fig. 1b, c)^46,47^. In addition, we observed a dramatic asymmetry in cross-hemisphere activity during turning, such that ipsilateral activity increased, and contralateral activity decreased, relative to the signal associated with forward walking (Fig. 1b, d). This observation, combined with previous studies^34,47^, suggested that interhemispheric differences in neural activity might correlate with the direction of turning, while symmetric increases in neural activity might correlate with forward movement. To test this hypothesis, we took advantage of the fact that the fly brain is almost perfectly bilaterally symmetric, allowing us to generate supercluster pairs that were spatially identical across the midline. Next, we conditioned behavior on the joint distribution of the neural activity for each left-right pair of superclusters (Fig. 1e-g; Extended Data Fig. 4).

Strikingly, every supercluster pair that correlated with walking followed a pattern in which the difference in neural activity across hemispheres correlated with angular velocity, while the sum of neural activity across hemispheres correlated with forward velocity (Fig. 1e-g; Extended Data Fig. 4). Thus, orthogonal changes in neural activity across the brain are associated with moving forward and turning.

### The symmetry break in network activity originates very early, in a small brain region

We next sought to define when and where this asymmetric pattern of interhemispheric differences first emerges relative to a change in angular velocity. We first implemented a bout triggered analysis, identifying every instance of a saccade, as defined by a peak in angular velocity >200 deg/sec, aligned these peaks in time, and averaged neural activity across all bouts (n=3786 bouts; Fig. 2a, b). The average saccade-triggered angular velocity across these bouts had a mean of zero until approximately 2 seconds before the turn, meaning that any interhemispheric differences in neural activity that emerged earlier than two seconds before the saccade must be related to the future turn, not to ongoing changes in behavior. Next, we empirically defined four temporal windows that preceded the turn (t=0) by up to 30 seconds, and computed the difference in activity for each left and right pair of superclusters, for all 3,786 saccadic turns (Fig. 2c). This analysis revealed that more than 15 seconds before a turn, no superclusters displayed interhemispheric differences. Strikingly, between 15 and 8 seconds before a turn, a single supercluster centered in the IPS displayed a large interhemispheric difference in neural activity that was dependent on the direction of the future saccade (Fig. 2c; Extended Data Fig. 5). A second supercluster in the visual system displayed a much weaker direction-dependent interhemispheric difference (Fig 2c; Extended Data Fig. 5). Between 8 and 0.4 seconds before a turn, these neural asymmetries expanded to include additional superclusters, with the majority of supercluster pairs displaying significant differences around the time of the turn (between 0.4 seconds before the turn and 0.4 seconds after the turn). Thus, the brain-wide asymmetry that exists at the time of a saccade emerges first between 15 and 8 seconds before the turn is executed within a specific brain region, the IPS.

**Fig. 2:**
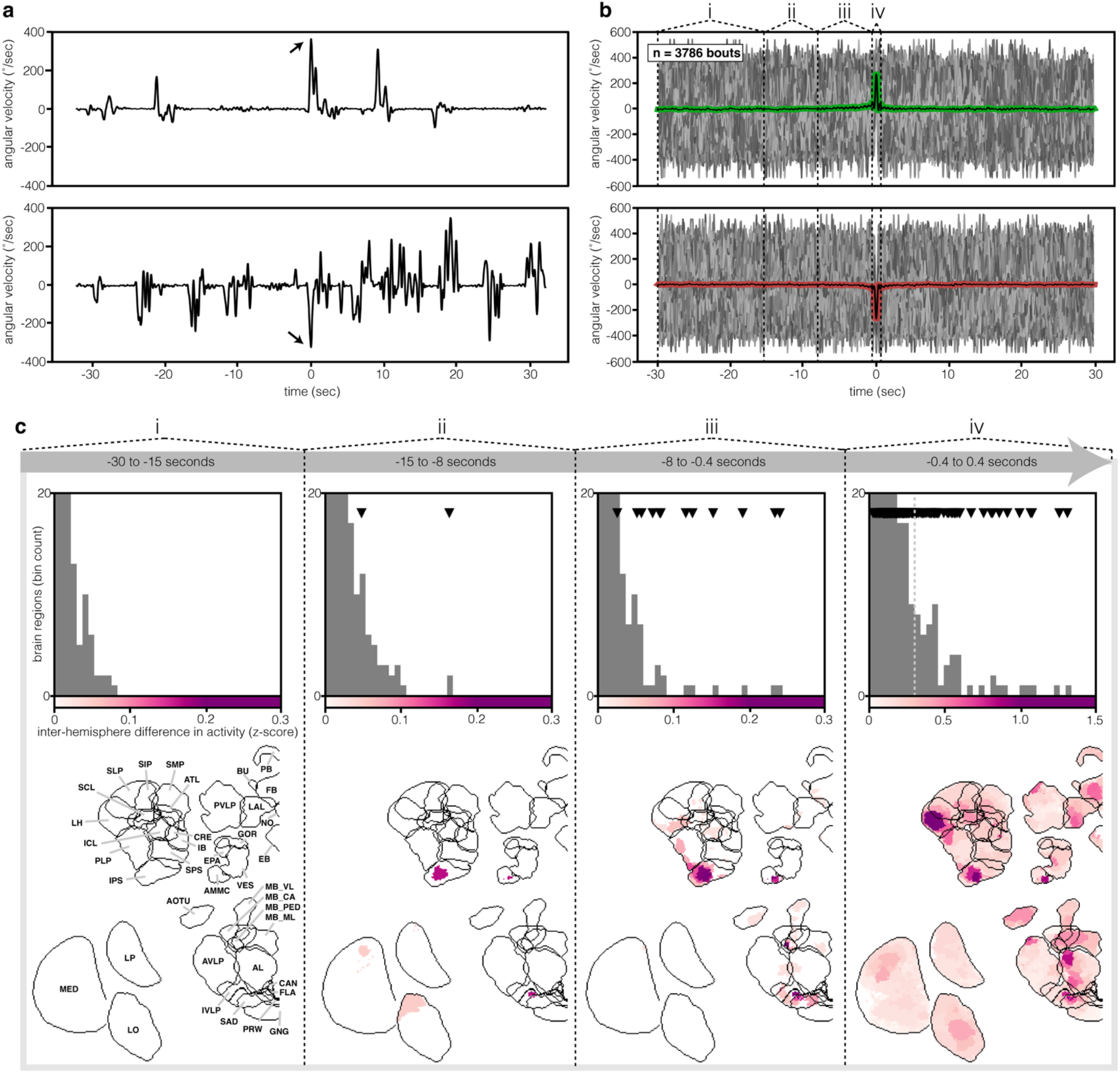
The turn-associated inter-hemispheric symmetry break originates very early in a small brain region and propagates across the brain. **a**, Two example behavioral traces, aligned at time zero relative to a peak in angular velocity for a left turn (top), or right turn (bottom). **b**, Overlay of all turn traces, aligned to peak angular velocity as in **a**, in grey scale. Turns were defined as having a peak angular velocity of >200deg/sec. Colored lines are the average of all left (top) and right (bottom) turns. n=3786 turns from across 9 flies. Dashed vertical lines and roman numerals correspond to time windows of interest used in **c. c**, Snapshots of neural activity within the four time-windows relative to a turn defined in **b**. Top histograms show the difference between neural activity in the left and right hemispheres for each supercluster. Upside-down triangles indicate regions that have a statistically significant difference (p<0.00004 based on Bonferroni corrected t-test with p<0.01 and 250 region comparisons). Exploded brain view, displaying regions that are colored by inter-hemispheric differences in activity (bottom). Only significant superclusters are colored.

To determine the precise timing of the emergence of inter-hemispheric asymmetries in neural activity relative to turning, we generated linear filters that relate neural activity to behavior across the entire dataset for each pair of superclusters (Fig. 3a,b and Methods). Using this approach, differences in the relationship between neural activity and behavior for superclusters on each side of the brain are revealed as differential weighting of the linear filter with respect to each turn direction. For example, when the fly turned left, the linear filter for a supercluster on the left side of the brain had a larger weight, and peaked earlier, than the filter for the corresponding voxel on the right side of the brain. We then computed the difference between these filters for each pair of superclusters in each hemisphere, when the fly was turning either left or right. Consistent with our bout-triggered analysis, the filter for the sub-region of the IPS diverged for left and right turns 15 seconds before the turn, while turn-selective signals emerged in the filters for many other brain regions much later, beginning approximately 3 seconds before the turn (Fig. 3). Within this 3 second period, regions involved in navigation (FB, PB, EB) and motor control (LAL, IPS) were recruited before the early visual system (MED, LO, LP, PVLP AVLP) and regions involved in mechanosensory processing (AMMC) (Fig. 3d,e). Thus, the early turning signal in the IPS emerges many seconds before most other regions of the brain display turn-dependent signals, consistent with this region playing a unique role in planning future motor actions.

**Fig. 3:**
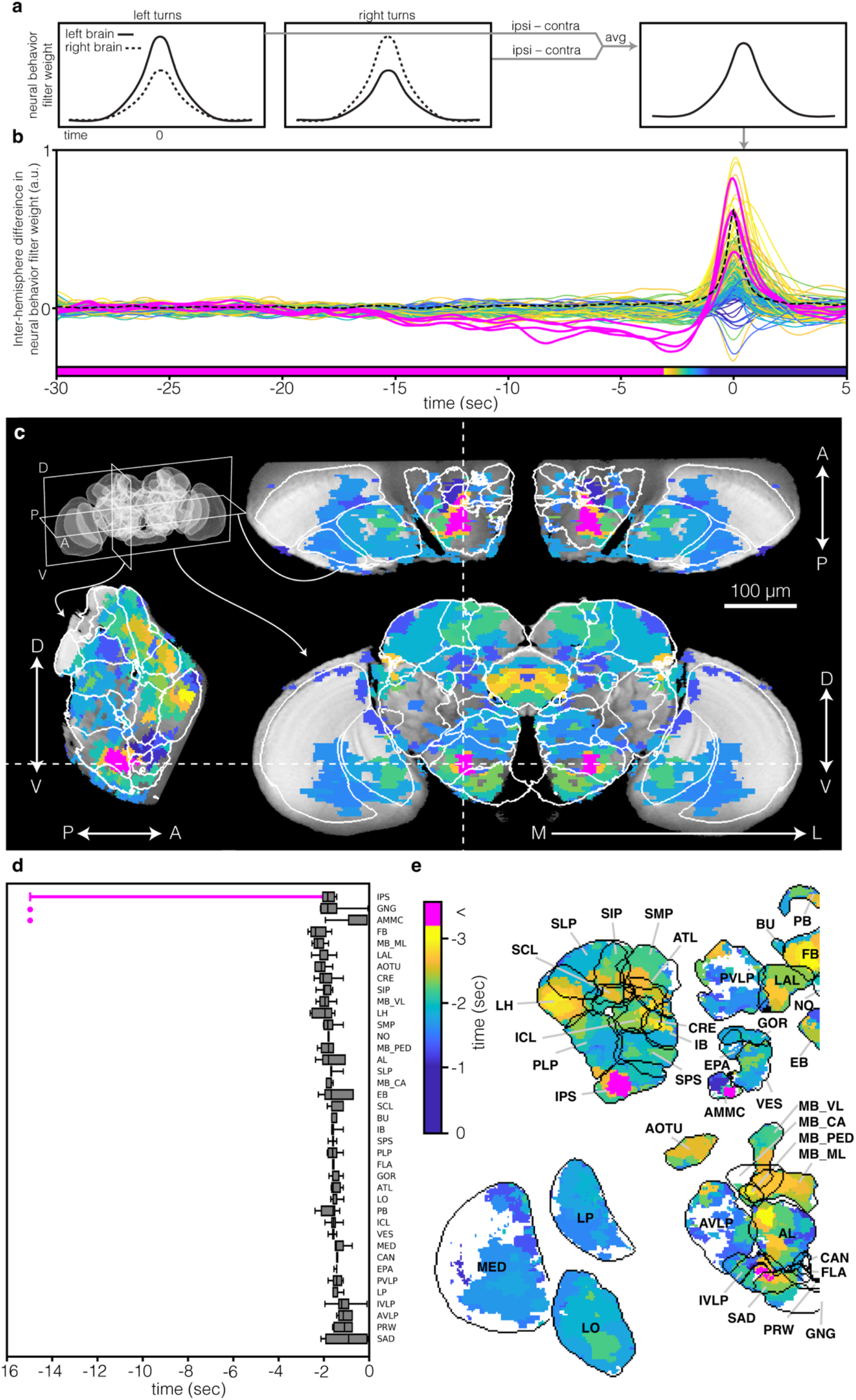
Temporal filters reveal the specific time of the inter-hemisphere symmetry break of each brain region. **a**, Schematic of temporal filter creation. For each matched pair of superclusters, the left and right turn filter is calculated separately for the left and right hemispheres. For both turn types, the contralateral filter is subtracted from the ipsilateral filter. Averaging these two filters gives the final single filter. **b**, Overlay of temporal filter from each supercluster. Color represents time the filter differs significantly from zero, meaning that the asymmetry has emerged (Methods). Magenta superclusters correspond to the supercluster identified in Figure 2, as well as three superclusters contiguous with the identified supercluster. Dashed black line corresponds to the autocorrelation of angular velocity, which serves as a useful comparison. **c**, Example slices through orthogonal planes of the brain colored by time of symmetry break. **d**, Box and whisker plots of distribution of symmetry break time for each anatomically defined brain region. Box center line indicates median, box limits indicate quartiles, whiskers indicate the 10^th^ and 90^th^ percentiles, dots indicate outliers. **e**, Exploded brain view colored by the time of symmetry break.

In this IPS sub-region, which we will now refer to as the pre-motor center, the level of neural activity diverged for left and right turns 15 seconds before the turn, held stable for more than 10 seconds, and then reversed sign to follow the brain-wide pattern of greater activity on the side of the brain that matches the turn direction (Fig. 3b, 4a). This pattern was consistent across flies (Extended Data Fig. 6). To directly visualize the relationship between neural activity and behavior, we created a phase plot of the neural-behavior space, both for single animals, and across the population, comparing the pre-motor center to a control brain region, the Lateral Horn (Extended Data Fig. 7). As expected, there was no structure along either the behavior axis or the neural activity axis up to 15 seconds before a saccade in either the pre-motor center or the Lateral Horn. However, between 15 and 8 seconds before the turn, while there was still no change along the behavioral axis, the neural axis in the pre-motor center (but not the Lateral Horn) bifurcated into separate regions of phase space, depending on the direction of the future turn, in both single flies and the population. Between 8 and 0.4 seconds, both behavior and neural axes were traversed in both the pre-motor center and the control, as travel along the neural axis increased, and behavioral change began. As the turn was executed, the trajectory made a large excursion along the behavior axis, while being held in a highly asymmetric neural state. Finally, we visualized the inter-hemisphere difference in neural activity across the joint angular and translational velocity distribution, for each time window (Fig. 4c, d). This analysis revealed that the asymmetric difference in neural activity in the pre-motor center extended across all rotational velocities of the animal, beginning 15 seconds before the turn, and was not modulated by forward velocity.

**Fig. 4:**
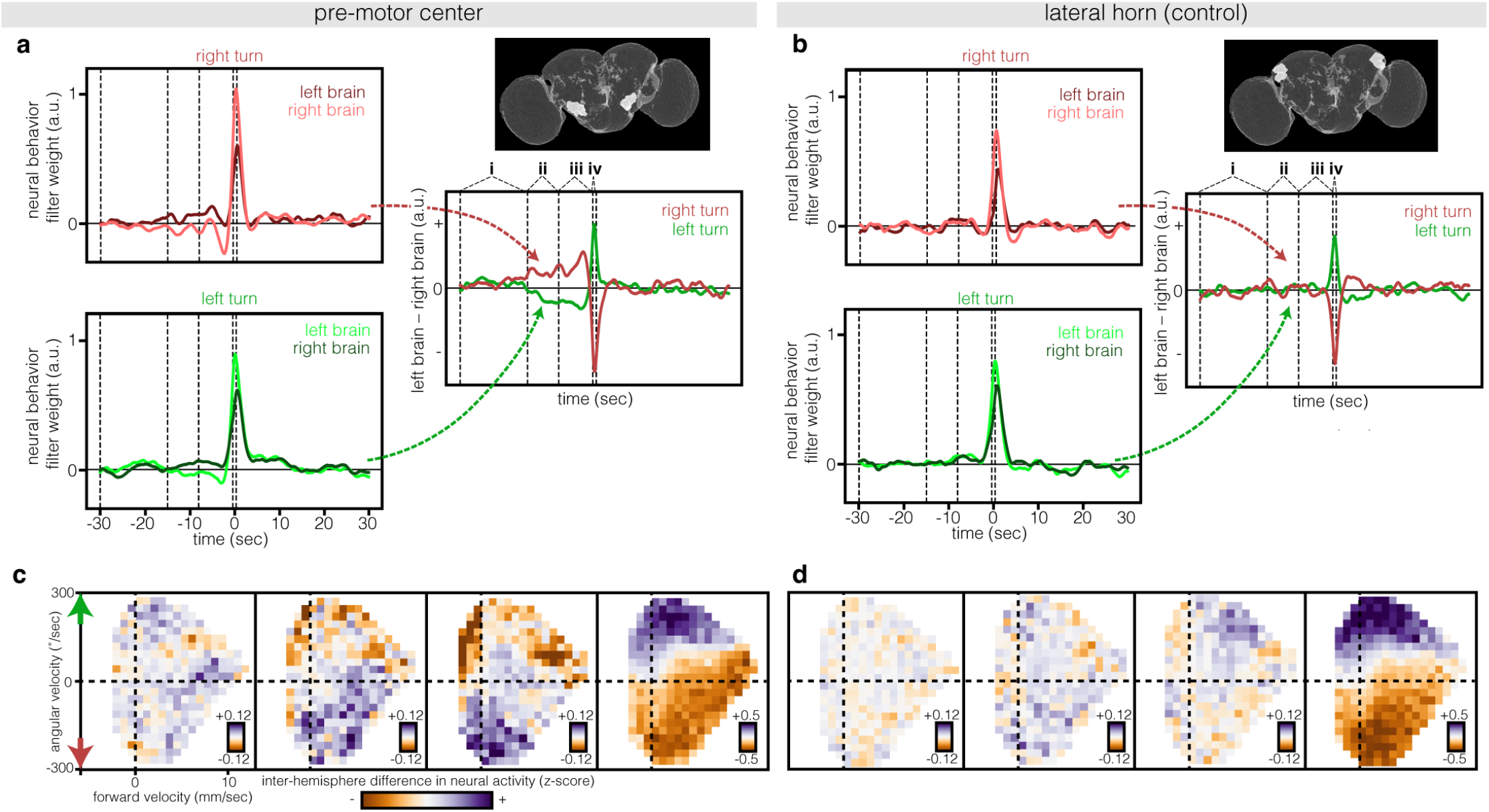
Neural activity in the pre-motor center precedes changes in velocity. **a**, linear filters for right and left turns, and left and right hemispheres, in the pre-motor center. Upper panel displays the pre-motor center in the context of the entire brain. Dashed vertical lines and roman numerals correspond to time windows of interest used in **c**,**d. b**, as in **a** but for a control region, the lateral horn **c**, Inter-hemispheric difference in neural activity conditioned on the joint distribution of forward and angular velocities, shown for each time window. **d**, as in **c**, but for the lateral horn.

We next examined whether the early signals in the pre-motor center corresponded to biases in the direction of future saccadic turns. First, we predicted the direction of individual turns, based on the neural activity in the pre-motor center from the preceding 15 to 8 second window, by computing the difference in neural activity across the pre-motor center region, setting a cutoff at 0. As a result, a positive value corresponds to a predicted turn in one direction, while a negative value corresponds to a predicted turn in the opposite direction. With this simple criterion, we could weakly but significantly predict the future turn direction by using neural activity in this brain region between 15 and 8 seconds before a turn across the entire dataset (54% correct, n=3786, p < 0.001, Fig. 5a, Extended Data Fig. 8, Methods). Conversely, predictions using signals from the Lateral Horn, or from the merged signal from all other superclusters except the pre-motor center failed to predict above chance (Fig. 5a). We reasoned that prediction accuracy is likely diminished by neural activity associated with ongoing behavior in the 15 to 8 second window, which could conflict with the predictive signal. To test this possibility, we selected for the subset of turns in which the fly was not moving during the prediction window (Extended Data Fig. 9). Under these conditions, prediction accuracy using pre-motor center activity between 15 and 8 seconds before a turn was increased substantially (66% correct, n=54, p < 0.001; Fig. 5a), while prediction accuracy derived from the Lateral Horn or the merged signal remained at chance levels (Fig. 5a). Finally, as expected, neural signals from the pre-motor center, the Lateral Horn, and the merged signal, were all predictive of turn direction when taken during the turn (Fig. 5a). Thus, early interhemispheric differences in the pre-motor center correspond to biases in the direction of future turns.

**Fig. 5:**
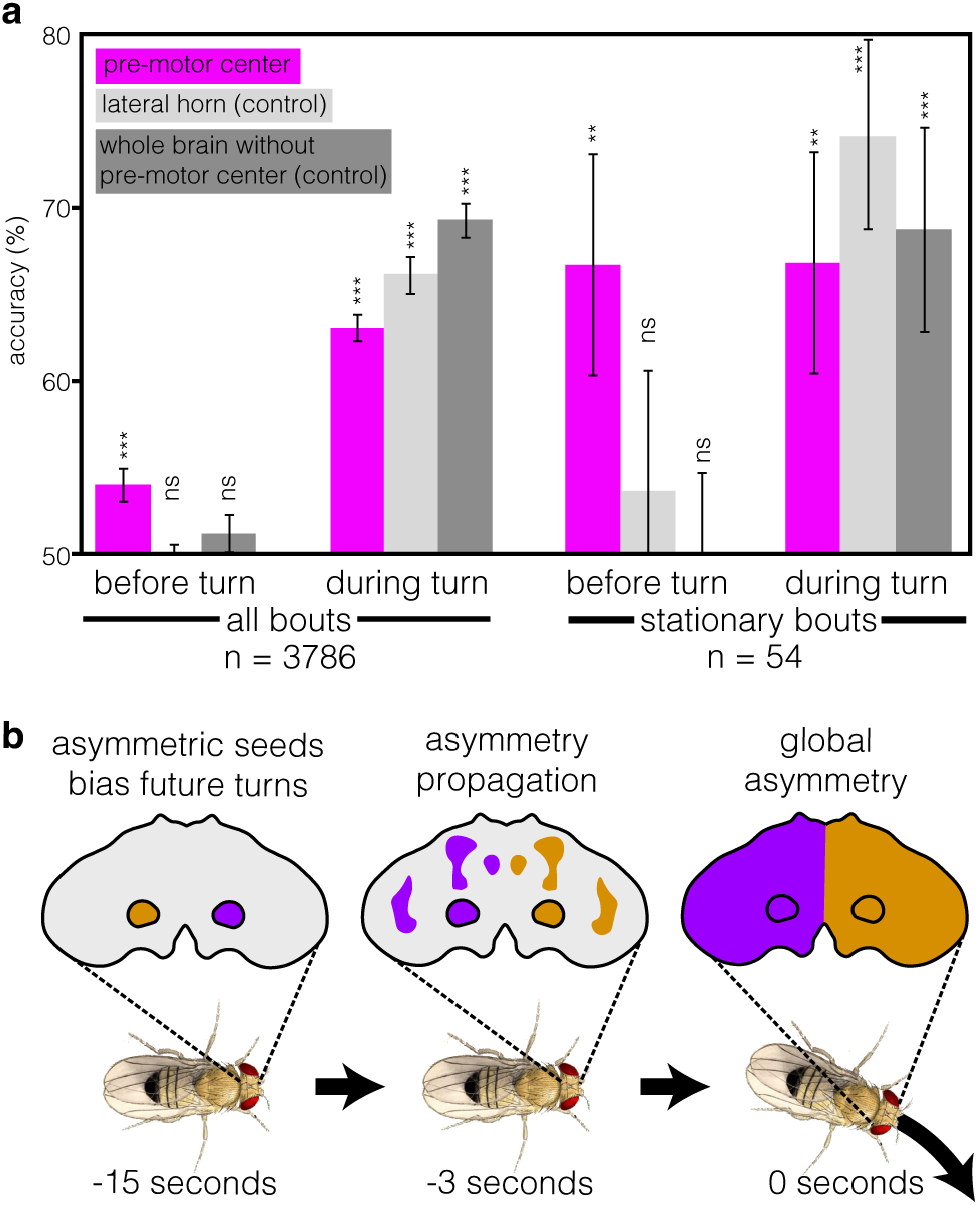
Neural activity in the pre-motor center biases future turn direction. **a**, Accuracy of predicting turn direction from neural activity. To generate these predictions, neural activity within the pre-motor center, the lateral horn (control), and all superclusters except those within the pre-motor center were used to predict turn direction. Predictions were generated using either the neural activity before (−15 to -8 seconds) or during (−0.4 to 0.4 seconds) a turn. Turn direction was predicted based on whether the difference in neural activity between matched interhemispheric pairs of superclusters, averaged across the time window, was above or below zero. These predictions were done for all turning bouts, as well as for a subset of turning bouts where the fly was stationary from -15 to -0.4 seconds before the turn. Error bars indicate the s.d. of a bootstrapped distribution. Comparing this distribution to the null hypothesis determined the p-value (Methods). *p < 0.05, **p < 0.01, ***p 0.001. **b**, Cartoon summarizing brain-wide spatiotemporal changes in neural activity that lead up to a turn. 15 seconds before a turn neural activity in the pre-motor region becomes asymmetric across the hemispheres, with an imbalance that biases future turns. As the time of the turn approaches, the asymmetry propagates across the brain, leading to a widespread asymmetry during the turn.

## Discussion

Although external sensory cues can instruct action selection, animals are nonetheless capable of generating specific movements in their absence. Here, we reveal neural correlates underlying voluntary action selection in the fly, and discover that neural activity in a bilaterally symmetric pre-motor center shapes a self-initiated turn more than 10 seconds in the future. Thus, this small animal is capable of planning future actions over a surprisingly long period of time. In the absence of sensory cues, flies stochastically choose between two intrinsically binary options: to turn left or to turn right. Our data demonstrate that the difference in neural activity between the ipsi- and contralateral pre-motor centers predicts this binary choice. This initial symmetry break between hemispheres also appears to catalyze the brain-wide asymmetry that emerges during turn execution (Fig. 5b).

The early activity in the pre-motor center is initially higher on the contralateral side of the brain, ramps over time, and then reverses to be higher on the ipsilateral side as the turn is executed. This pattern is strikingly similar to the lateralized readiness potential seen in complex vertebrates^53,54^. That this early activity is only partially predictive of the future direction of turning suggests that at early stages, the animal has not yet committed to a specific action, maintaining flexibility to allow additional internal signals to inform the choice of turn direction. We speculate that the activity within the pre-motor center represents a neural correlate of volition in the fly, setting the stage for future work that will dissect the mechanistic basis of this cognitive process at the cellular and circuit level.

## Extended Data Figures

**Extended Data Fig. 1:**
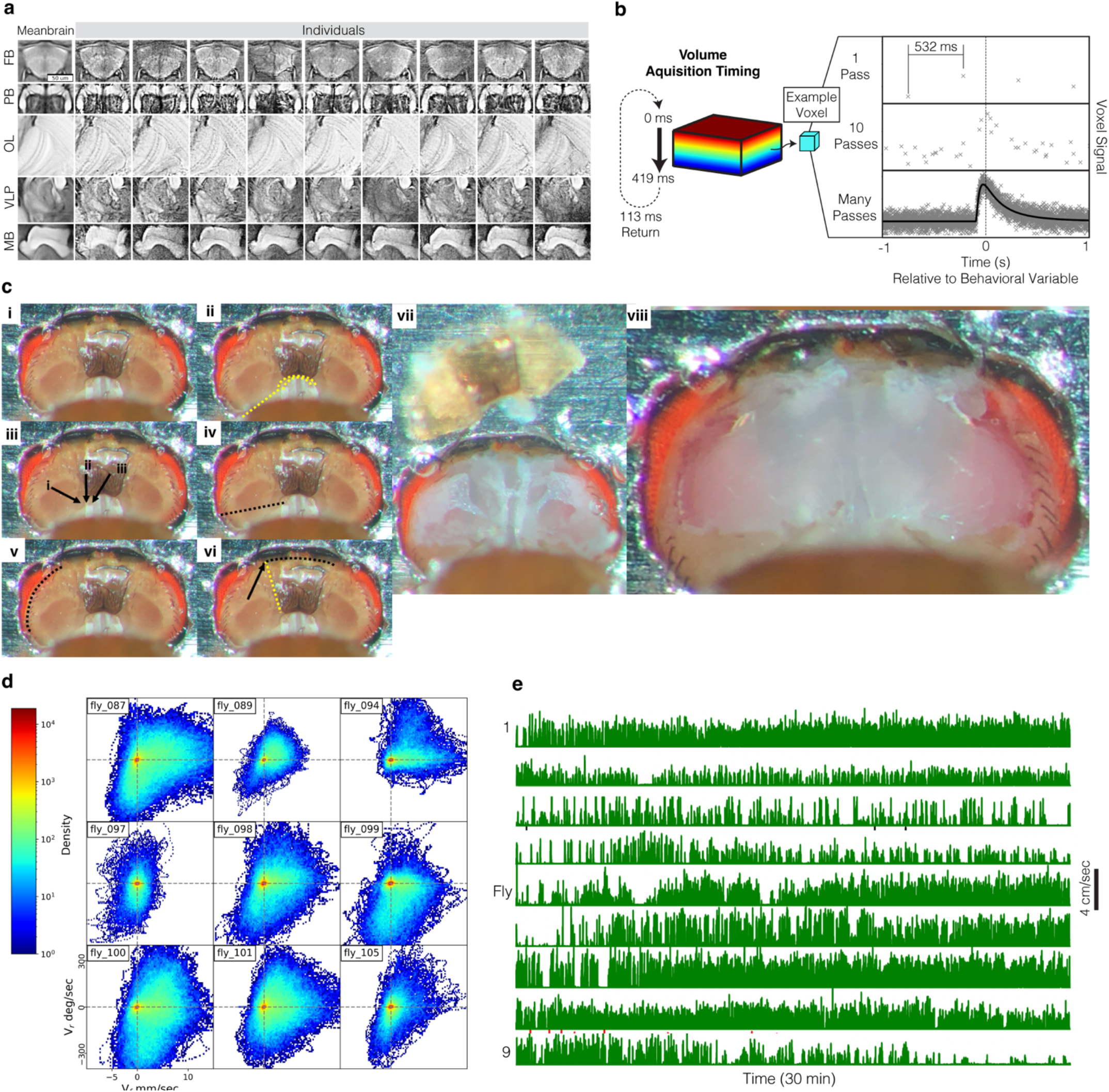
Whole-brain imaging and data alignment in walking *Drosophila*. **a**, Qualitative comparison of alignment quality. The average of all 9 brains (the “meanbrain”) is compared with anatomical scans from each individual. Five representative regions of the brain were cropped to the same coordinates after alignment. **b**, Schematic illustration of the logic for obtaining high temporal resolution filters from low temporal resolution data. The time of measurement for each voxel within the volume is precisely tracked. As each measurement of neural activity (defined by the scan time) is not synchronized to behavior, each imaging pass is offset from a chosen behavioral event (like a saccadic turn), meaning that while single scan would generate a poor estimate of the underlying linear filter relating neural activity to behavior, many scans at different temporal offsets allow high temporal resolution filter to be computed. **c**, Overview of the sequence of dissections needed to expose the entire brain. (i) Each fly was glued into a metal shim. (ii) Placement of first cuts with dissection needle. (iii) Three cuticle boundaries near the neck. (iv) Cut along the black line. (v) Cut along the eye border. (vi) Yellow line denotes a strong cuticular structure. Continue cutting along the black line, breaking through yellow line. (vii) Remove cuticle. (viii) Remove trachea and fat. **d**, 2D histograms of forward (Vf) and rotational (Vr) velocity for the nine flies in this study. **e**, Temporal traces of walking behavior derived from ball movement in any axis across the entire 30 minute session for each fly.

**Extended Data Fig. 2:**
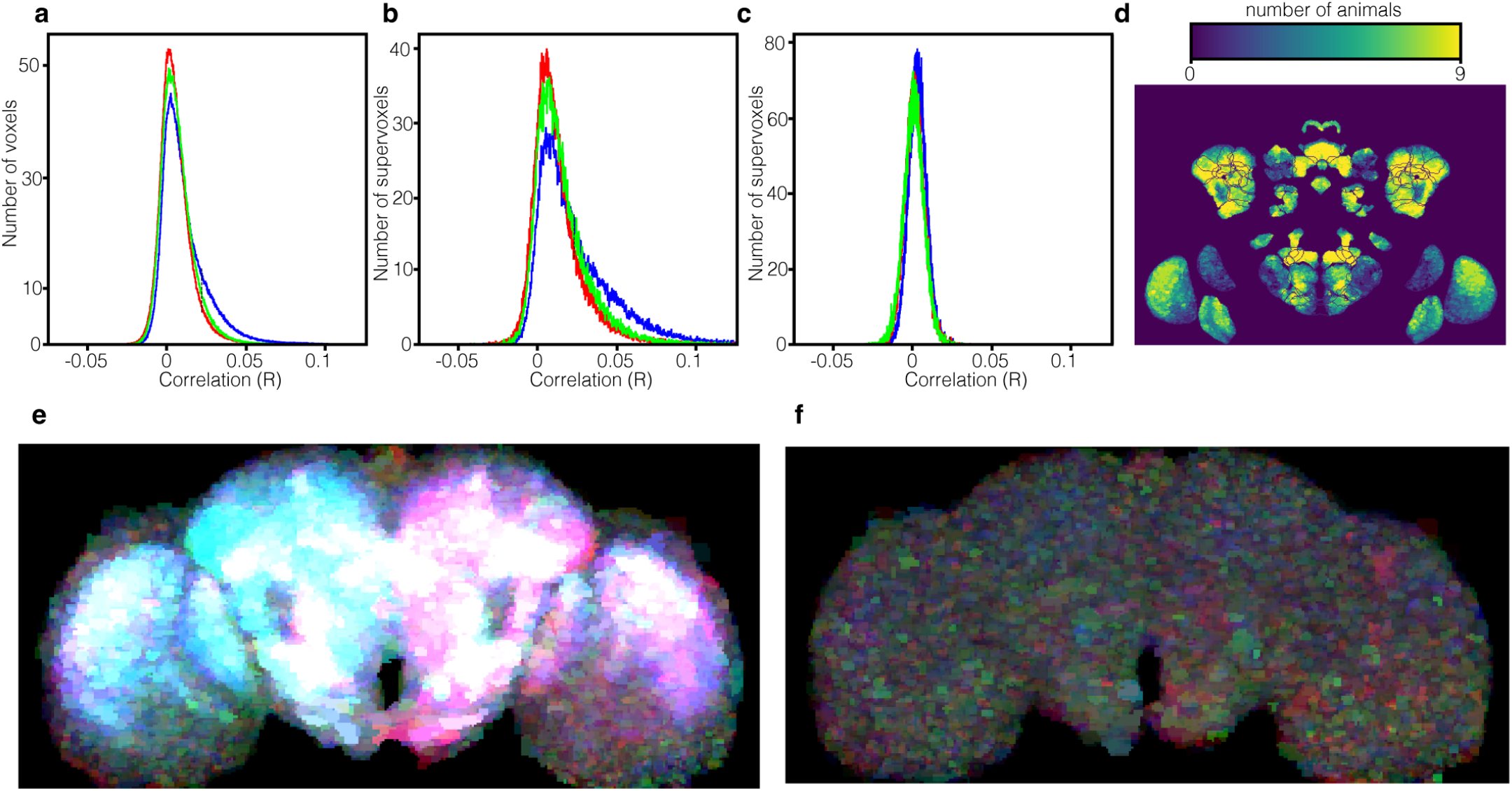
Brain-wide correlations measured with single voxels versus supervoxels using both GCaMP and tdTomato. **a**, Distribution of single voxel correlations with three velocity components (blue: translational velocity; red: angular velocity turning right; green: angular velocity turning left). **b**, As in **a**, but calculated using supervoxels. Note that the correlations between neural activity and behavior have increased due to the increased SNR. **c**, As in **b**, but calculating the correlations using the tdTomato signal instead of GCaMP6f. The structural marker had no significant correlations with behavior, ruling out contributions from brain motion. **d**, Brain-wide map of neural activity, displaying the number of flies that had a correlation with velocity above an r-value of 0.05 for each voxel. This analysis demonstrates that all flies make similar contributions to the dataset. **e**, Maximum intensity projection through all z-slices, overlaying each voxels correlation with each of the three velocities (in blue, green and red). **f**, As in **e**, but using the correlations with each of the three velocities calculated using the tdTomato signal. The absence of signal in this panel indicates that motion artifacts were undetectably small.

**Extended Data Fig. 3:**
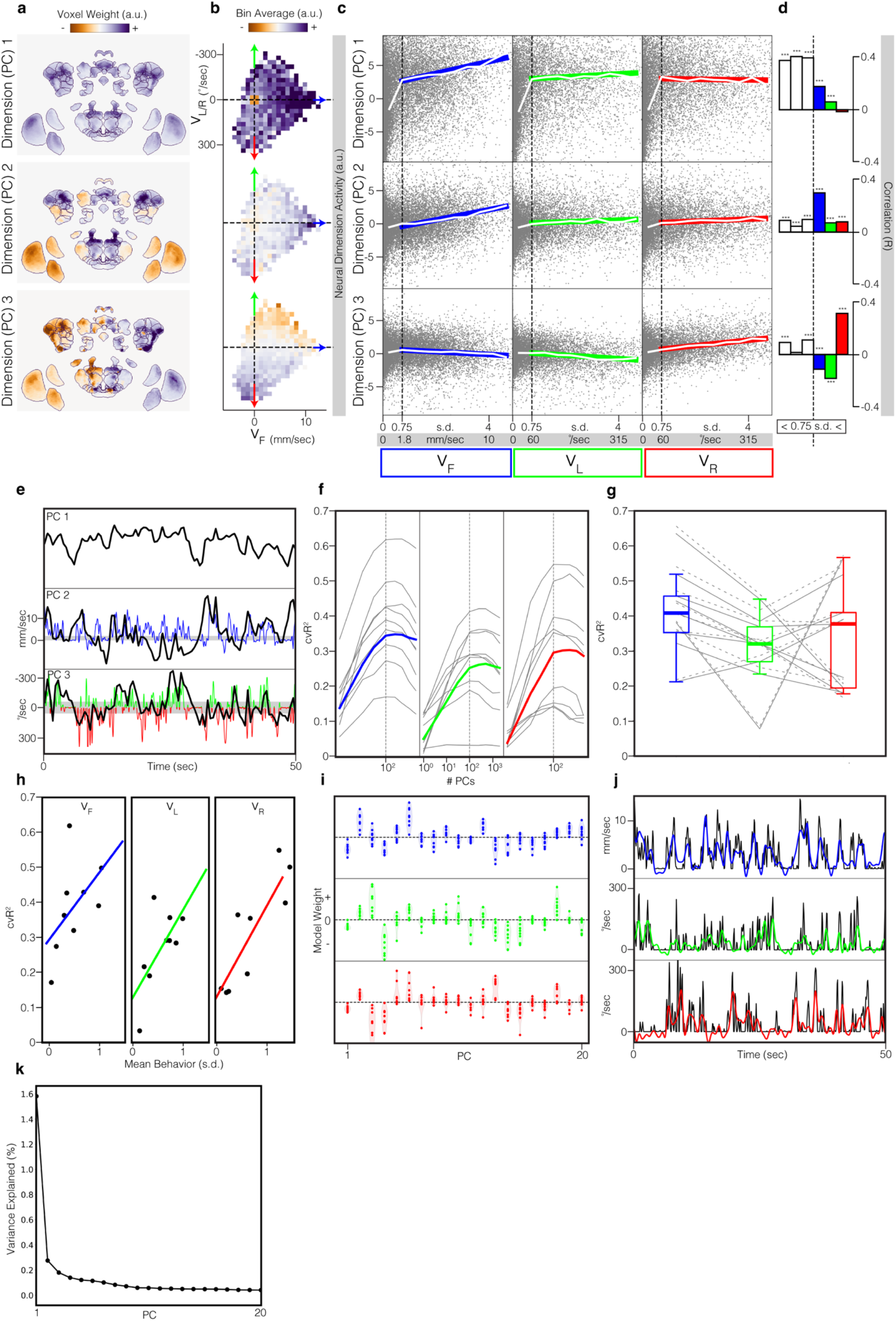
Low dimensions of neural activity are structured and predictive of velocity space. **a**, The first three principal components of the whole-brain neural dataset (n = 9 flies). Maximum intensity projections are colored by voxel weights for each principal component. **b**, Neural data projected onto each principal component and binned in 2D velocity space. We note that the small skew in the angular velocity distribution reflects a small rotational asymmetry in placement of each fly on the ball. **c**, 1D velocity spaces versus principal component values. Grey dots are single timepoints of a collected neural volume. Vertical dashed black line represents the chosen threshold between moving and not-moving. Colored lines denote the linear regression of data above this threshold. White lines are mean bin values, with a width of 0.5 s.d.. Vertical white lines represent ± 1 SEM, but are too small to be visible. **d**, Quantification of data in **c**. Correlations were measured independently for below and above the movement threshold. *p < 0.05, **p < 0.01, ***p 0.001. **e**, Example trace of first 3 PCs and behavior. Black lines are PCs, blue is forward velocity, and red and green are rotational velocities. PCs are overlayed with relevant behaviors. Gray shading shows 0.75 s.d. threshold used in previous panel. **f**, Cross-validated R2 prediction accuracy on test data for linear models independent fit to predict velocity components. For each velocity component, separate models were fit using different numbers of principal components as input features (x-axis). Curving gray lines are individual flies. Colored lines are the integrated superfly. Vertical dashed gray lines represent the number of components used in subsequent panels. **g**, Cross-validated R2 prediction accuracy for a model with 100 input principal components. Solid gray lines are individual flies predicted using the principal components from the integrated superfly dataset, while the dashed gray lines are individual flies fit using their respective individual principal components. Box plots represent first quartile, median, and third quartile, while whiskers are 1.5 times the interquartile-range. **h**, Same data as solid gray lines in **f**, but compared with the mean amount of each velocity component seen in each individual fly. Colored lines are linear regression. **i**, Model weights fit for the first 20 principal components. Each dot is an individual fly. **j**, Actual (black) versus predicted (color) velocity traces. Principal components and behavior were interpolated to 10 Hz for this visualization. **k**, Neural variance explained by the first 20 principal components. This variance explained plot is strongly shaped by the shot noise inherent in resonant scanning.

**Extended Data Fig. 4:**
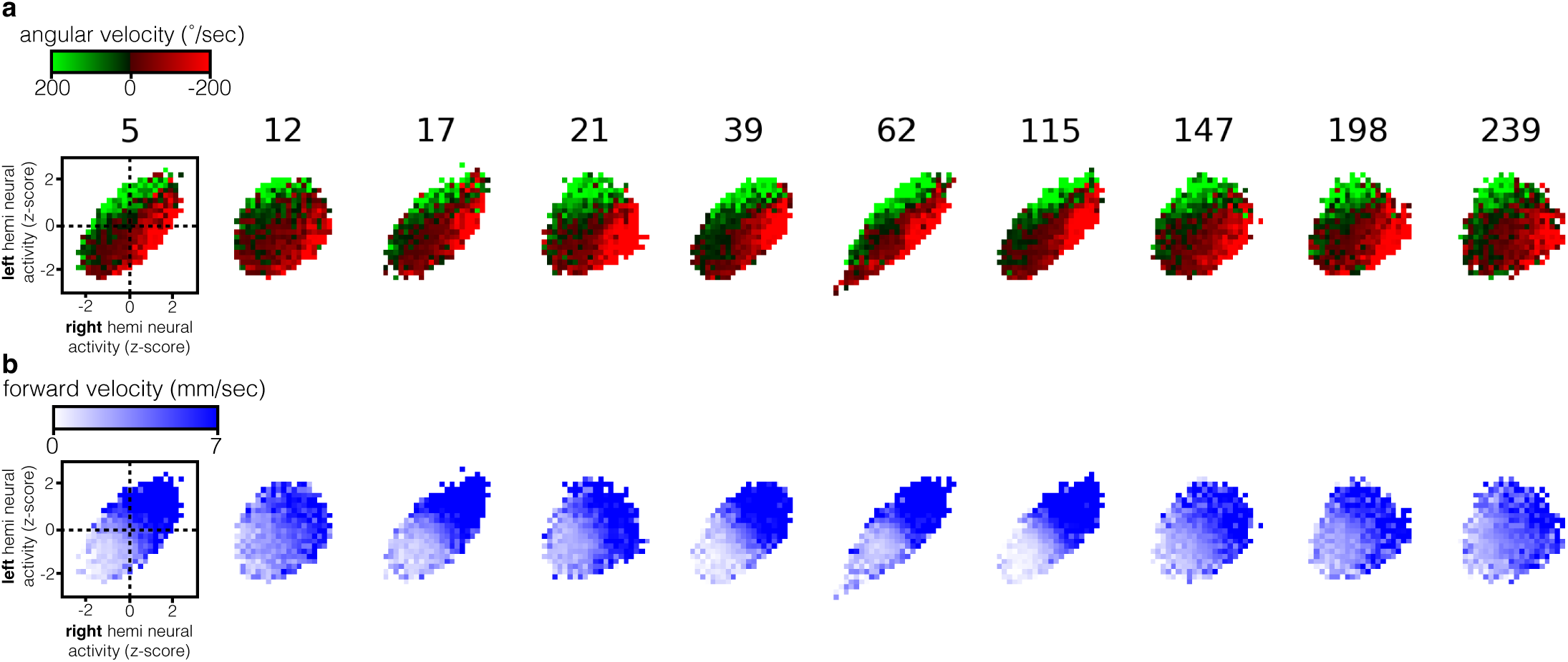
Representation of the relationship between angular velocity and translational velocity for additional inter-hemispheric pairs of supervoxels. Extended from Figure 1 e-g. Top: angular velocity conditioned on joint neural activity across the two hemispheres. Bottom: as in top row, but with forward velocity. These super-voxels display similar relationships to behavior as those displayed in Fig. 1, demonstrating the brain-wide nature of this representation of behavioral correlates.

**Extended Data Fig. 5:**
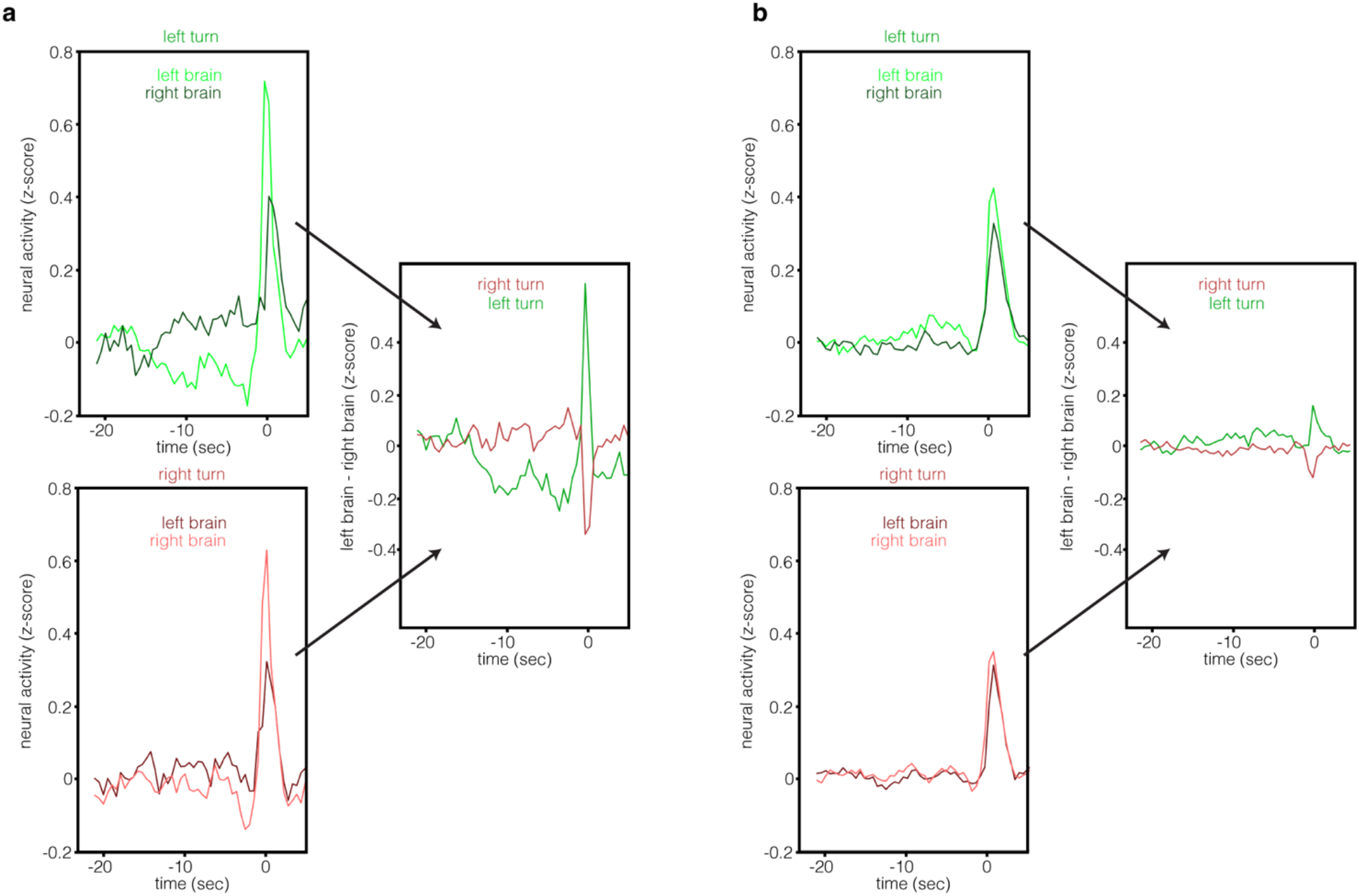
Bout-triggered neural activity of superclusters significant within the -15 to -8 second time window before a turn. **a**, supercluster 77, located in the Inferior Posterior Slope (IPS). Neural activity is averaged across all left and right bouts. **b**, same, but for supercluster 106, located in the Lobula (LO)

**Extended Data Fig. 6:**
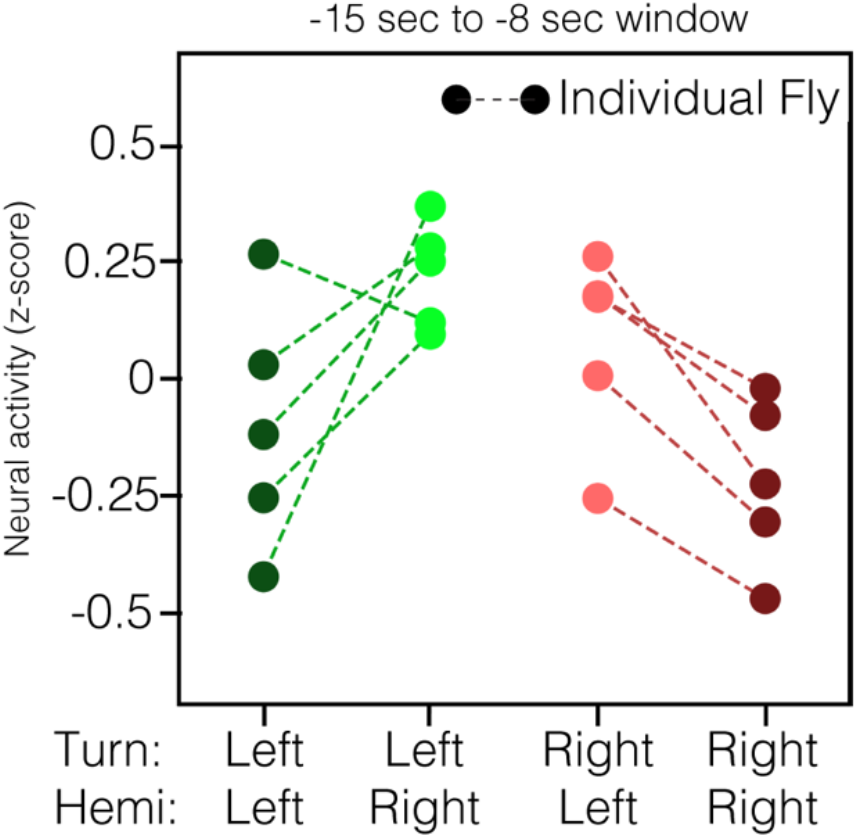
Average neural activity of the bout-triggered neural average (all bouts) of the pre-motor center within the -15 to -8 seconds time window for individual animals. Flies that did not have at least 50 turn bouts in each direction were excluded from this analysis. During this early window, neural activity is higher in the right pre-motor center than the left pre-motor center for left turns, while neural activity is higher in the left pre-motor center than the right pre-motor center for right turns (p=0.008 with Wilcoxon rank-sum test).

**Extended Data Fig 7:**
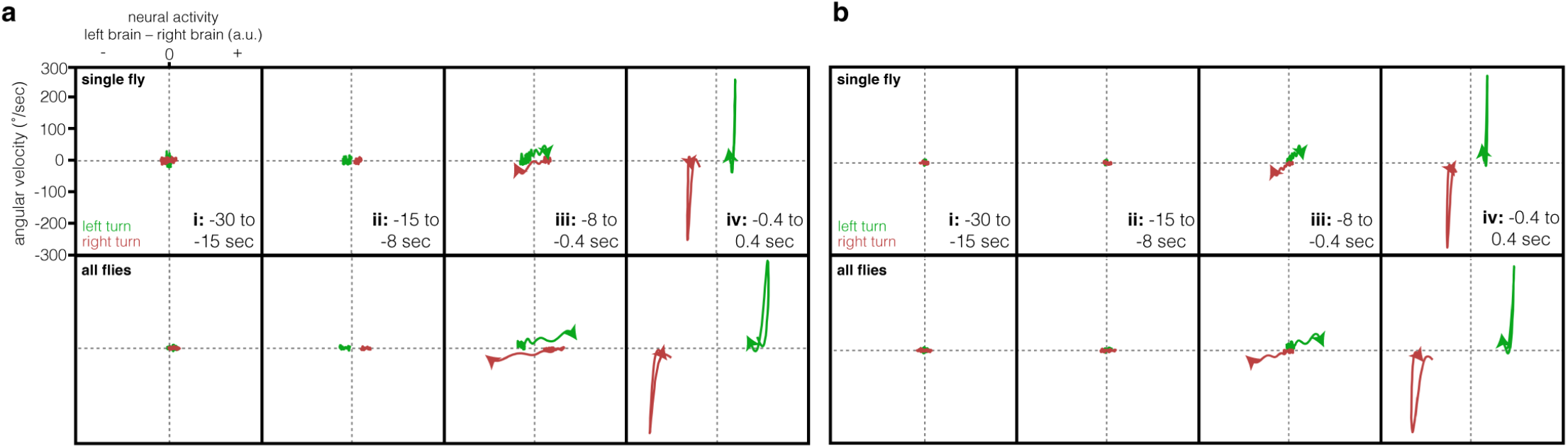
Phase space plots show the relationship between neural activity and behavior within each time window relative to a turn. Difference in neural activity between the left and right hemispheres is plotted (x-axis) versus angular velocity (y-axis). Temporal windows are as defined in Figure 2 and Figure 4, and indicated within each panel. Top row; a single example fly. Bottom row; average of all flies. **a**, neural activity from the pre-motor center. **b**, neural activity from the lateral horn (control).

**Extended Data Fig 8:**
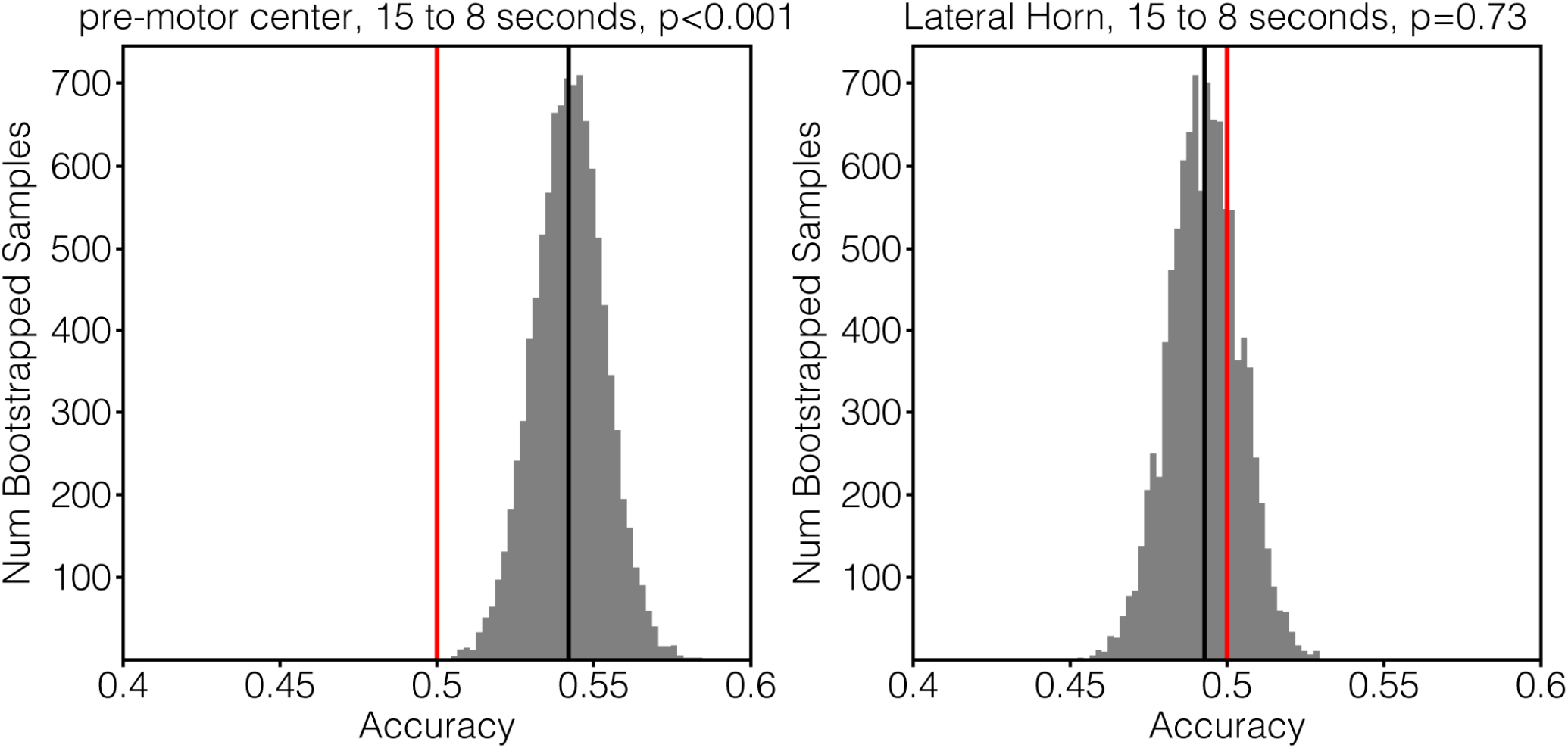
Bootstrap distributions of turn direction predictions. Related to Figure 5a and Methods. Histogram shows distribution of the 10,000 bootstrapped samples. Vertical black line is the mean accuracy across samples. Vertical red line is null hypothesis of no predictability. P-value is the fraction of samples that have an accuracy below chance.

**Extended Data Fig. 9:**
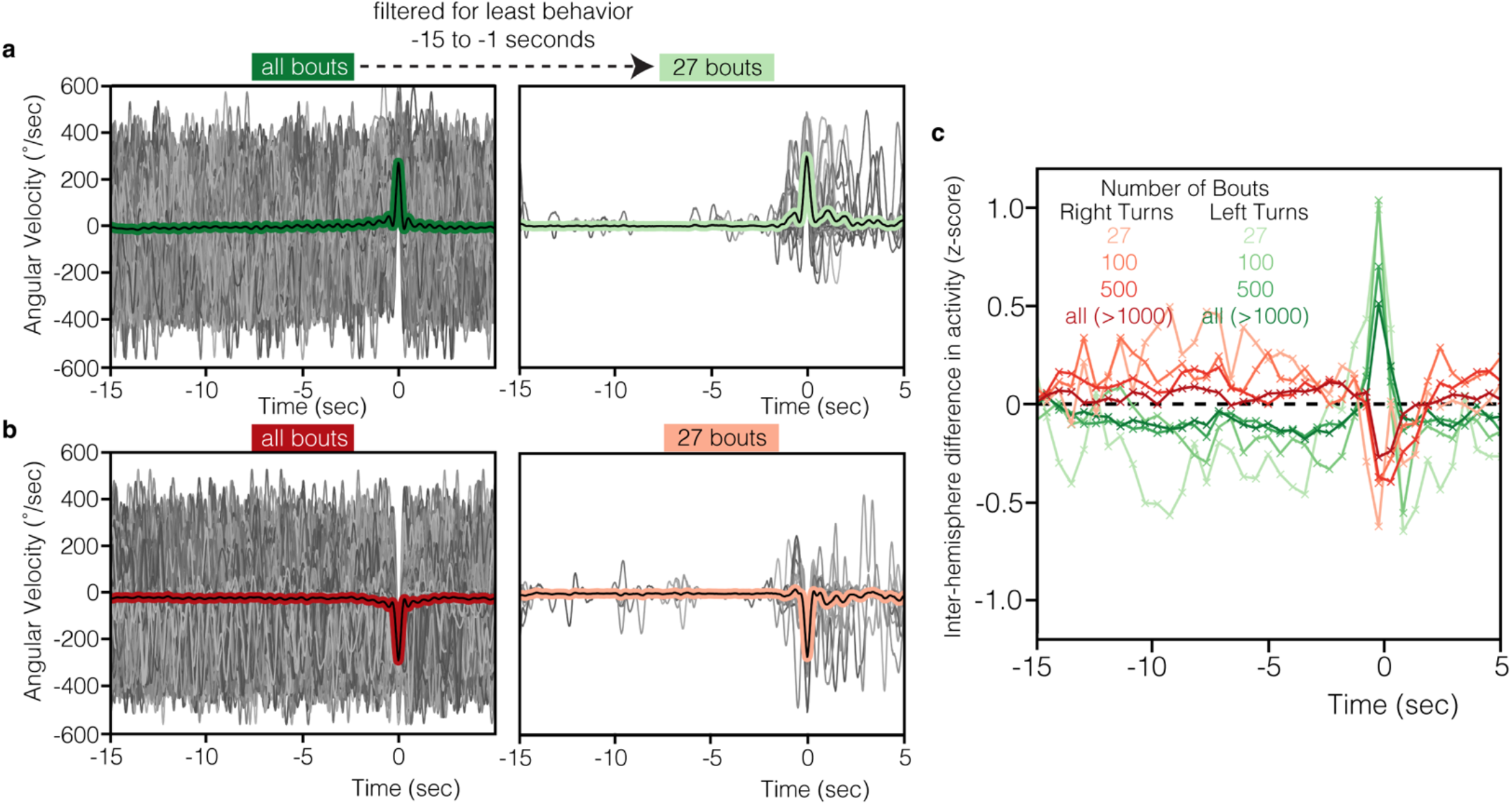
Filtering turn bouts for different levels of behavior before the turn increases aniticipatory inter-hemispheric neural differences in the pre-motor center. **a**, Overlay of all left turns. Turns were defined as having a peak angular velocity of >200deg/sec. Green line is the average of all turn bouts. Right panel is an overlay of the 27 left turns in which the flies were virtually stationary within the time window of -15 to -1 seconds. **b**, As in **a**, except for right turns. **c**, Bout-triggered average of neural activity within the Inferior Posterior Slope. Neural activity is left hemisphere minus right hemisphere. Red shades are right turns, green shades are left turns. Bouts have been sorted by amount of behavior within -15 to -1 second time window, such that the “27” bouts are the 27 bouts with least early behavior, and similarly the “100” bouts are the 100 bouts with the least early behavior, etc.

## Methods

### Experimental Models

*Drosophila melanogaster* were of the genotype *w+/w+;UAS-myr::tdTomato/UAS-GCaMP6f;nSyb-Gal4/+*. Flies were raised on molasses medium at 25 °C with a 12/12-h light/dark cycle. Flies were housed in mixed male/female vials of 10-20 individuals. 3-4 days post-eclosion females were used for imaging.

### Mounting and Dissection

Each fly was anesthetized on a chilled Peltier plate with a thermally coupled custom holder. Each immobilized fly was carefully fitted into a custom mount consisting of 3D-printed plastic and a custom cut steel shim to tightly nestle the head and thorax. To fix the fly to the mount, UV-curable glue was placed and cured on the dorsal region of the face between the eyes, and on the thorax. A saline solution was added to the dish for dissection (103 mM NaCl, 3 mM KCl, 5 mM TES, 1 mM NaH_2_PO_4_, 4 mM MgCl_2_, 1.5 mM CaCl_2_, 10 mM trehalose, 10 mM glucose, 7 mM sucrose, and 26 mM NaHCO_3_). Using a tungsten needle the posterior head cuticle was carefully cut and removed to reveal the whole brain (Extended Data Fig. 1). Dissection forceps were used to remove fat and trachea.

### Two-Photon Imaging

Flies were imaged using a resonant scanning Bruker Ultima IV system with a piezo drive and a Leica 20× HCX APO 1.0 NA water immersion objective lens. GCaMP6f and tdTomato were simultaneously excited with a Chameleon Vision II femtosecond laser (Coherent) at 920 nm. A 525/50 nm filter and a 595/50 nm filter were used to collect signals from GCaMP6f and tdTomato. Photons were detected simultaneously using two GaAsP-type photomultiplier tubes. The exposed fly brain was perfused with carbogen-bubbled (95% O_2_, 5% CO_2_) saline solution (same as above) heated to 30°C with an in-line heater. For the 30 min functional scan, volumes were collected at a resolution of 2.6 × 2.6 × 5 μm (256 voxels x 128 voxels x 49 slices, XYZ), resulting in an approximate volume rate of 1.8Hz. Scans were bidirectional along the X axis. For the immediately subsequent anatomical scan, spatial dimensions were adjusted to 0.6 × 0.6 × 1μm (1024 voxels x 512 voxels x 241 slices, XYZ), and 100 volumes were collected. In this orientation, all regions of the brain were visible, except for the laminas in each optic lobe, which are occluded by the eye, and a portion of the Gnathal Ganglion, which is occluded by the esophagus.

### Behavior Tracking

During imaging, the head-fixed fly performed spontaneous bouts of walking on a painted, air-suspended foam ball (9 mm diameter, LAST-A-FOAM FR4615). The ball was imaged at 50 Hz with a Flea FL3-U3-13E4M-C sensor and Edmund Optics 100 mm C Series Fixed Focal Length Lens. An IR LED directed with optic fibers was used to illuminate the ball. Frames were processed using Fictrac to calculate the animal’s walking velocity^48^. Before all subsequent analysis, forward and rotational velocities were smoothed with a Savitzky-Golay filter of window length 500 ms and a polynomial of order 3.

### Data Preprocessing

Brain volumes were first motion-corrected using ANTs^49,50^; the tdTomato channel was time-averaged across the 30 min recording and each tdTomato volume was warped (affine and non-linear) to the mean. Each volume’s warp parameters were then applied to the GCaMP6f channel. Then, each voxel was independently corrected for bleaching as well as other slow temporal trends by subtracting a temporally smoothed signal from the raw trace (smooth signal produced by gaussian filter of 2 minute sigma, truncated at 1 sigma). Finally, each voxel in the GCaMP6f recording was Z-scored. This preprocessing was all done on individual animals before volumetric alignment and concatenation.

### Data Alignment

Data alignment was performed as in Brezovec et al^45^. Each individual’s anatomical scan was created from 100 collected volumes of tdTomato signal. These 100 volumes were first averaged, then each volume was warped (affine and non-linear) to this mean using ANTs. These aligned volumes were then averaged, creating the final anatomical scan for the individual. These anatomical scans were then all registered into a common brain space: the Functional Drosophila Atlas (FDA). Each individual’s functional scan was affine aligned to their anatomical scan using the tdTomato channel and applying the transforms to the GCaMP6f channel. Then, each anatomical scan was aligned (affine and non-linear) to the FDA using the tdTomato channel, and the transforms were applied to the GCaMP6f channel. The FDA contains anatomical region delineations^51,52.^

### Agglomerative Clustering Supercluster Creation

Individual voxels were spatially aggregated into supervoxels, and finally superclusters, to reduce the number of features for computational tractability and to boost SNR as follows. After GCaMP6f data was preprocessed and warped into the meanbrain space, individual flies were temporally concatenated to create a single large matrix of (x, y, z, t; 256, 128, 49, 30456) (Fig. 1a). Then, single voxels were merged independently for each z-slice via agglomerative clustering with Ward linkage and a connectivity constraint (only spatially neighboring voxels could be merged), resulting in 2000 supervoxels per z-slice. Next, we performed a second round of agglomerative clustering. Here we are merging across z-slices and while constraining for bilaterally symmetric superclusters to facilitate inter-hemisphere analyses. Here we took advantage of the principal components calculated from the supervoxels (Methods). These PCs can reveal spatial structure that may be less resolvable by simply using the raw temporal traces, and may therefore create clusters with the best agreement in neural signals across many dimensions of neural activity. We used the first 100 spatial components since this is where our linear models peaked in behavior predictability (Extended Data Fig. 3f). Because we use these superclusters to compare neural signals in matching regions across the two hemispheres, we enforced symmetry by mirroring and averaging the absolute value of each spatial principal component across the hemisphere-midline. We then used the same agglomerative clustering algorithm used to create the supervoxels (ward linkage with a connectivity constraint). We determined the number of superclusters by sweeping a range (from 100 to 10,000) and visualized the resulting brain-wide correlation and cross-correlation maps. We selected 500 clusters (250 in each hemisphere) as optimal since it minimized the number of superclusters while avoiding mixing clusters that were distinct in terms of correlation and temporal cross-correlation properties.

### Principal Component Analysis and Linear Modeling

Principal components were calculated using the entire dataset with supervoxels (supervoxel, z, t; 2000, 49, 30456). The matrix was reshaped as (feature, t; 98000, 30456), the covariance matrix was calculated, and an eigendecomposition was performed producing eigenvalues and eigenvectors. This was repeated for individual flies as well (Extended Data Fig. 3g). Linear models were fit using principal components as features and a single behavioral variable as output. The number of features were swept to find the value that maximizes prediction accuracy (Extended Data Fig. 3f). Data was split into training and test sets and five-fold cross-validation was used to calculate the prediction accuracy on the held out test set (R^2^). A ridge penalty was employed to regularize the model and prevent overfitting.

### Linear Filter Analysis and Definition of Symmetry Break Point

Given a matrix of timestamps for each slice of neural data acquisition across the superfly (z, vol_num; 49, 30456), windows of interpolation of behavior variables were created centered at each slice and volume number, with 20 ms steps extending 5 sec before and after. This resulted in a behavior matrix of (z, vol_num, interp_window; 49, 30456, 500). This matrix was then weighted by the neural activity for each supervoxel to produce a filter matching the interpolation window. This resulted in the equivalent of a cross-correlation filter for each supervoxel and for each behavior variable. We noticed these filters had a power peak in the frequency spectrum at the volume imaging rate (1.8 Hz), which we removed with a notch filter. Next, we deconvolved the GCaMP6f kinetics (Methods). Inter-hemisphere symmetry breaks were defined as the first timepoint before the filter peak that the derivative of the inter-hemisphere difference passed a set threshold (0.001 filter-weight / 20 milliseconds). Superclusters were excluded if their filter peaks were too weak (<0.1 filter-weight), or if the filters for left and right turns had a Pearson correlation below 0.5 R.

### GCaMP6f Deconvolution

The temporal dynamics were deconvolved from both the cross-correlation filters and the whole-brain spatiotemporal filters as follows. An experimental measurement of the GCaMP6f impulse response was used (100 ms to peak and 250 ms wide at half-max), taken from *Drosophila* early visual system neurons Tm3 and Mi1^55^. The measured kinetics were modeled by the formula ΔF/F = (1-e-t/4)*(-e-t/8)^56^. This was expanded into a Toeplitz matrix and the deconvolved filters are given by the least-squares solution to y = Xb, where y is the measured convolved filter, X is the Toeplitz matrix, and b is the unknown deconvolved filter^57^.

### Prediction of Turn Direction

All turn bouts across the 9 flies were identified by peaks in angular velocity >200deg/sec (n=3786). The neural activity trace from each supercluster for each individual bout (temporally centered on the bout peak) was collected into a list. For each bout, the neural activity trace from the right brain was subtracted from the neural activity trace of the left brain (for either the pre-motor center, the Lateral Horn, or the whole-brain except the pre-motor center). The neural trace was averaged over the temporal window of interest (either 15 to 8 seconds before the bout, or -0.4 to 0.4 seconds flanking the bout). One round of bootstrapping involved selecting 1,000 left turn bouts and 1,000 right turn bouts at random and with replacement. The accuracy of a single round was calculated as (num_correct_left/1,000 + num_correct_right/1,000)/2, where num_correct_left is the number of left bouts with a neural activity above zero, and num_correct_right is the number of right bouts with a neural activity below zero. Note the sign was flipped for the pre-motor center early time prediction, since left turns correspond to neural activity below zero, and right turns correspond to neural activity above zero (Fig. 4a). The accuracy was calculated over 10,000 bootstrapped samples. The null hypothesis is an accuracy of 0.5, so the one-tailed p-value was determined as the number of bootstrapped samples that were under 0.5 divided by 10,000.

## Author Contributions

L.E.B. and T.R.C. conceived the project. L.E.B. collected experiments and analyzed the data. A.B.B. registered neural data with SynthMorph. S.D. provided modeling direction and feedback. L.E.B. and T.R.C. wrote the manuscript. T.R.C. advised throughout the project.

## Acknowledgements

We would like to thank the Dickinson lab, in particular Will Dickson, for providing us with treadmill balls, as well as the Maimon lab, in particular Jazz Weisman, for direction on building our fly mounting apparatus. We would also like to thank members of the Clandinin Lab, as well as Albert Lin for feedback on the manuscript. This work was supported by a National Science Foundation Graduate Research Fellowship (LEB), as well as grants from the NIH (R01EY022628 (TRC), R01EY022638 (TRC), 5U19NS104655 (TRC and SD), 3R01NS11006003 (TRC)), the Stanford Vision Core Grant 5P30EY02687704 (TRC), and the Simons Foundation (TRC). TRC is a Chan-Zuckerberg BioHub Investigator.

## Declaration of Interests

The authors declare no competing interests.

## Data Availability

Microscopy (structural and functional neural recordings) and behavioral data have been deposited at Dryad and are publicly available as of the date of publication.

## Code Availability

All original code has been deposited at Github and is publicly available as of the date of publication. https://github.com/ClandininLab/brezovec_volition_2023

## Notes

### Competing Interest Statement

The authors have declared no competing interest.

